# Microstimulation in the primary visual cortex: activity patterns and their relation to visual responses and evoked saccades

**DOI:** 10.1101/2020.05.03.072322

**Authors:** R. Oz, H. Edelman-Klapper, S. Nivinsky-Margalit, H. Slovin

## Abstract

Intra cortical microstimulation (ICMS) in the primary visual cortex (V1) can generate the visual perception of phosphenes and evoke saccades directed to the stimulated location in the retinotopic map. Although ICMS is widely used, little is known about the evoked spatio-temporal patterns of neural activity and their relation to neural responses evoked by visual stimuli or saccade generation. To investigate this, we combined ICMS with Voltage Sensitive Dye Imaging in V1 of behaving monkeys and measured neural activity at high spatial (meso-scale) and temporal resolution. Small visual stimuli and ICMS evoked population activity spreading over few mm that propagated to extrastriate areas. The population responses evoked by ICMS showed faster dynamics and different spatial propagation patterns. Neural activity was higher in trials w/saccades compared with trials w/o saccades. In conclusion, our results uncover the spatio-temporal patterns evoked by ICMS and their relation to visual processing and saccade generation.

## Introduction

Intra-cortical Microstimulation (ICMS) is a well-established technique to artificially stimulate the brain that enabled to uncover the organization and function of the cortical areas as well at their connections to other brain structures. Importantly, ICMS enabled to study the causal link between cortical areas to sensation, movement and cognition (reviews: Clark, Armstrong, & Moore, 2011; Cohen & Newsome, 2004; Histed, Ni, & Maunsell, 2013; Tehovnik & Slocum, 2007). While the recently developed optical stimulation techniques offer improved spatial resolution and cell type specific activation or suppression (El-Shamayleh and Horwitz 2019; Kim et al. 2017), ICMS is still extensively applied. Moreover, ICMS is currently the preferred stimulation technique in research and applications involving the development of cortical prosthesis and brain-machine interfaces (Bosking et al. 2017; Lewis et al. 2015; Richardson et al. 2019; Schmidt et al. 1996)

Electrical stimulation and ICMS in the primary visual cortex (V1), has been proposed as a tool to restore vision in blind humans, with accumulating efforts for about half a century (Brindley and Lewin 1968; Dobelle and Mladejovsky 1974; Lewis et al. 2015; Schmidt et al. 1996; Tehovnik et al. 2009). However, despite the extensive research, the spatio-temporal patterns of the neural activity evoked by ICMS remains poorly understood. Previous studies have reported that the amount of current injected through an microelectrode to activate a cortical neuron is directly proportional to its square distance from the electrode tip, implying that ICMS generates neural activity within a small area around the electrode (Tehovnik 1996; Tehovnik et al. 2004). However, Histed et al. (2009) reported on sparse neural activation distributed across a wide area, where the activity in remote regions relative to the stimulating electrode, was suggested to be mediated by axons passing nearby the stimulated region. In addition, the propagation of activity generated by electrical stimulation to higher areas is under debate. Several studies reported that microstimulation can propagate one hierarchical level, only one synapse away, while cortical areas two synapses away are suppressed (Brock et al. 2013; Logothetis et al. 2010; Sultan et al. 2011), however, it was recently suggested that neural activity can propagate from V1 to areas that are more than one hierarchical level higher (Fehérvári et al. 2015; Fehérvári and Yagi 2016; Klink et al. 2017).

The sensory experiences evoked by electrical stimulation were reported to have some similarities with normal sensory stimulation (Histed et al. 2013). In particular, electrical stimulation in V1 of humans were shown to produce the visual sensation of a small point of light, termed as phosphene (Brindley and Lewin 1968; Clark et al. 2011; Foerster 1929; Penfield and Perot 1963). ICMS in monkeys’ V1 produces behavioral effects that were consistent with phosphene induction (Clark et al. 2011; Tehovnik and Slocum 2007). The position of a phosphene in the visual field is corresponding to the stimulated region within the retinotopic map in V1, i.e the location of the stimulated receptive fields (RFs) in the visual field (Bradley et al. 2005; Brindley and Lewin 1968; Penfield and Perot 1963; Schmidt et al. 1996). Another well-established effect of ICMS in V1 is the generation of saccades directed to the stimulated site in the retinotopic map of V1 which is also the RF location of the stimulated neurons (Doty 1965; Tehovnik and Slocum 2013).

Although ICMS has been used extensively, little is known about the spatio-temporal patterns of neural activity in V1 evoked by ICMS and their relation to responses evoked by visual stimuli or to saccade generation. To investigate this topic, we performed ICMS in V1 of two monkeys while they were performing a fixation task and recorded population activities using voltage-sensitive dye imaging (VSDI, Shoham et al., 1999; Slovin, Arieli, Hildesheim, & Grinvald, 2002). In addition, we imaged the population responses evoke by small visual stimuli while the animals were maintaining fixation and compared the time course of the evoked population response and their spatial patterns under the different stimulation conditions. Finally, we compared V1 responses in ICMS trials with saccades to the stimulated RFs and compared them to trials w/o saccades to the stimulated RFs.

## Methods

### Intra-cortical microstimulation

A microelectrode was inserted through the artificial dura (Arieli et al. 2002) and targeted to the upper layers in V1. Biphasic square pulses were delivered through a standard tungsten microelectrode (FHC, Bowdoin, ME, USA) using a microstimulation box (linear biphasic stimulus isolator, BAK electronics, BSI-1A). Each biphasic pulse was comprised from a cathodal (0.2 ms) pulse followed by an anodal (0.2 ms) pulse. We stimulated the brain with current amplitude of 60-100 μA; Stimulation frequency: 333 or 500 Hz; Stimulation length: 15-240 ms (3-80 pulses). The output current from the microstimulation box was verified as voltage measurement across a 100 KΩ resistor located between the animal and the microstimulation box. The ICMS parameters we used were previously shown to be highly effective in evoking neural activity in visual cortices, evoking Phosphenes and induce saccades towards the RF of the stimulated neurons as well as affecting behavioral responses (Doty, 1965; Tehovnik, Slocum, & Schiller, 2003; Bradley et al., 2005; Bartlett et al. 2005; for review: Tehovnuk and Slocum 2013).

### Behavioral paradigm with ICMS or visual stimulation

Two adult male monkeys, *Macaca fascicularis* (monkey L and A, 6 and 7 years old respectively) were trained on a simple fixation task that was combined with ICMS or presentation of a small visual stimulus. The animals were required to maintain fixation throughout the entire trial. A trial started when the animal acquired fixation on a small (0.1 deg) white fixation point displayed on a uniform gray background. The animal had to maintain fixation within a small fixation window for a random time interval (3-4 s) and then a short train of ICMS was applied. Trials ended 1 s after ICMS onset when the fixation point was turned off and the animal was rewarded with a drop of juice for correct trials. Because ICMS can evoked saccades, in these sessions, the animals were allowed to make saccades within the fixation window. Trials with ICMS (30-50% of total trials) were interleaved with fixation-alone trials (no ICMS; blank trials). The blank trials were used to remove the heartbeat artifact and photo-bleaching of the VSD signal (see below). In a different set of experiments, the same two animals were presented with a small Gabor (wavelength (λ) - 0.25 deg; σ - 0.125 deg, phase - 0 deg; Figure 1B) while maintaining fixation. The visual stimulus was displayed for 200 ms and its position in the visual field was varied across different imaging sessions (eccentricities 0.75-3.5°), thus activating different regions within the imaged area.

**Figure 1:**
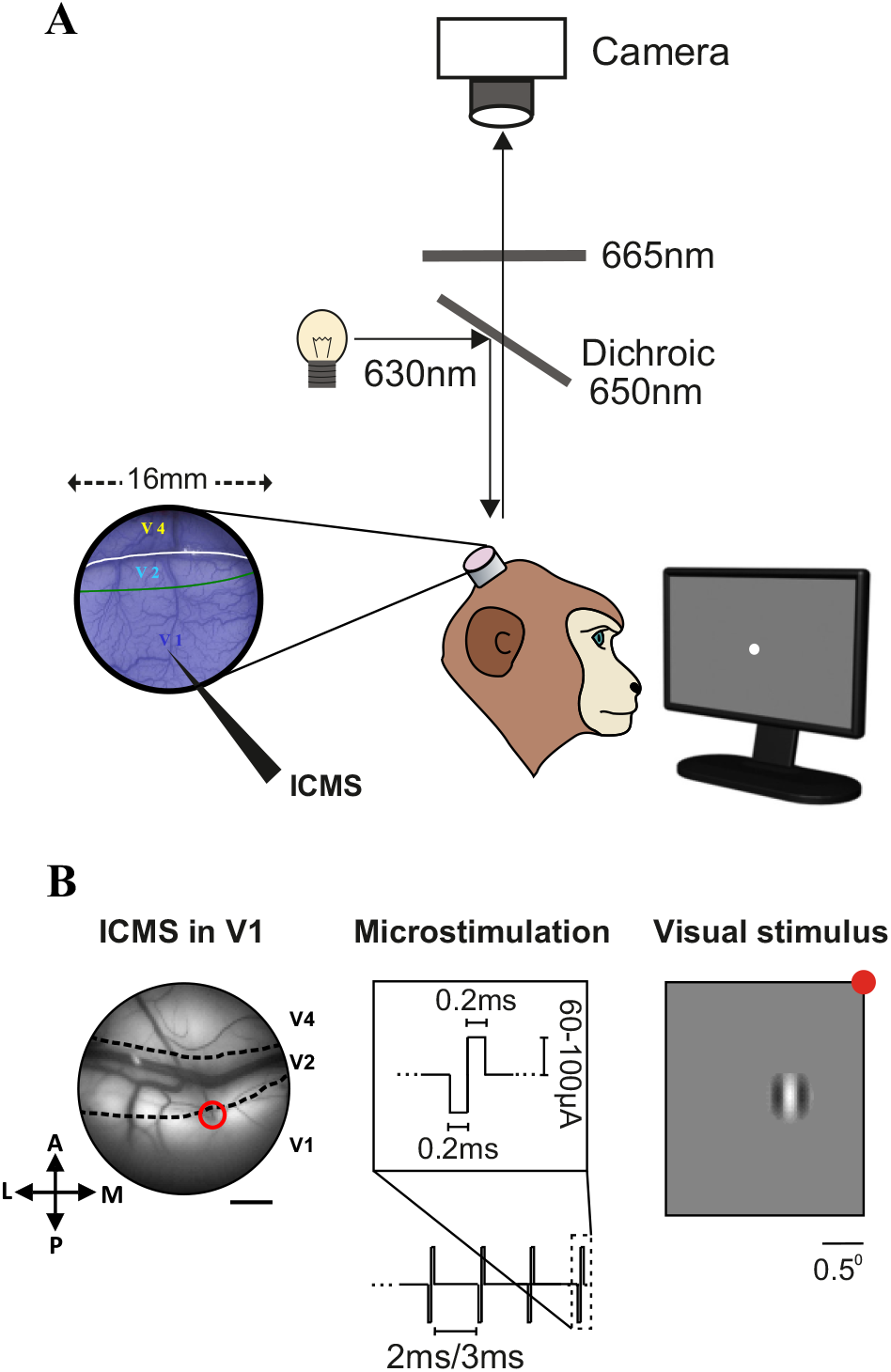
Illustration of the experimental setup and stimulation. **A.** Schematic illustration of the VSDI setup in behaving monkeys combined with ICMS in V1. The blue image on the left shows the imaged chambers after staining with the blue voltage-sensitive dyes (see Methods). The black elongated arrow in V1 denotes the microelectrode. **B.** ICMS was applied in V1 of two fixating monkeys where ICMS trials were interleaved with blank trials (no stimulation). Left: image of the blood vessels patterns in V1, V2 and V4 areas with a microelectrode inserted (marked with a red circle) through the artificial dura into the upper layers of V1 (monkey A). Scale bar is 2 mm. Middle: ICMS parameters: biphasic square pulses were delivered through the microelectrode, each pulse was comprised from a cathodal (0.2 ms) pulse followed by an anodal (0.2 ms) pulse. Range of current amplitude: 60-100 μA; Stimulation frequency: 333, 500 Hz; Stimulation length: 15-240 ms (3-80 pulses). Right: Visual stimulus: a small Gabor patch (2σ=0.25°; 100% contrast) was presented over the screen while the animal was maintaining fixation on a small fixation point (red point, for illustration). On different imaging sessions the visual stimulus appeared at different eccentricities (see Methods). Scale bar is 0.5°.

### Surgical procedure and voltage-sensitive dye imaging

The surgical procedure and VSD staining have been reported in detail previously (Arieli et al. 2002; Slovin et al. 2002). All experimental procedures were approved by the Animal Care and Use Guidelines Committee of Bar-Ilan University and supervised by the Israeli authorities for animal experiments and conformed to the NIH guidelines. Briefly, the monkeys were anesthetized, ventilated, and an intravenous catheter was inserted. A head holder and two cranial windows (25 mm, i.d.) were bilaterally placed over the primary visual cortices and cemented to the cranium with dental acrylic cement. After craniotomy, the dura mater was removed, exposing the visual cortex. A thin, transparent artificial dura of silicone was implanted over the visual cortex. Appropriate analgesics and antibiotics were given during surgery and postoperatively. The center of the imaged V1 areas were 1-3 ° below the horizontal meridian and 1-2° from the vertical meridian for both monkeys, the size of the exposed imaged area covered ~3–4°×4–5° of the visual field, at the reported eccentricities. We stained the cortex with RH-1691 or RH-1838 voltage-sensitive dyes (VSD; supplied by Optical Imaging). The procedure for applying VSDs to macaque cortex is described in detail by Slovin et al. (2002). VSDI was performed using the Micam Ultima system based on a sensitive fast camera with up to 10 KHz sampling rate. We used a sampling rate of 10 ms/frame with a spatial resolution of 10,000 pixels where each pixel summed the activity from an area of 100×100 μm^2^ (for ICMS sessions) or 170×170 μm^2^ (for visual stimulation sessions), every pixel summing the neural activity mostly from the upper 400-600 μm of the cortex. The exposed cortex was illuminated by an epi-illumination stage with appropriate excitation filter (peak transmission 630 nm, width at half-height 10 nm) and a dichroic mirror (DRLP 650), both from Omega Optical. To collect the fluorescence and reject stray excitation light, a barrier postfilter was placed above the dichroic mirror (RG 665, Schott).

### Experimental setup, visual stimulation and DAQ

Two linked personal computers managed visual stimulation, data acquisition, and controlled the monkey’s behavior. We used a combination of imaging software (MicamUltima) and the NIMH-CORTEX software package. The behavior PC was equipped with a PCI-DAS 1602/12 card to control the behavioral task and data acquisition. The protocol of data acquisition in VSDI was described previously (Slovin et al. 2002). To remove the heartbeat artifact, we triggered the VSDI data acquisition on the animal’s heartbeat signal (see information in VSD data analysis, and Slovin et al., 2002). Visual stimuli were presented on a 21 inch CRT Mitsubishi monitor at a refresh rate of 85 Hz. The monitor was located 100 cm from the monkey’s eyes.

### Eye position recording and saccade detection

Eye position was monitored by an infrared eye tracker (Dr. Bouis Device, Kalsruhe, Germany), sampled at 1 kHz and recorded at 250 or 500 Hz. The animals were required to maintain fixation within a small fixation window, however, as previously reported ICMS can evoke saccades towards the stimulated location in the retinotopic map of V1, which is also the location of the stimulated RFs in the visual field (Doty 1965; Tehovnik et al. 2003; Tehovnik and Slocum 2013). We therefore allowed the animals to make small saccades towards the RF location of the stimulated neurons, within the fixation window (although they were trained to maintain fixation). To detect the saccades on each trial, we implemented an algorithm for saccades and microsaccades (MSs) detection on the monkeys’ eye position data (Engbert and Mergenthaler, 2006). The algorithm could detect saccadic eye movements and MSs (larger than 0.1°) at high reliability, as well as their amplitude, direction and latencies. Previous studies from our group have already reported the correlation between saccades, MSs and the VSD signal in V1 and V2 (Gilad et al. 2014; Meirovithz et al. 2012).

### VSD data analysis

VSDI data were obtained from a total of 33 VSDI imaging sessions from two hemispheres of two monkeys. ICMS analysis was done on 5 and 20 imaging sessions from monkeys L and A respectively. The analysis on ICMS evoked saccades and VSD signal were performed on a total of 7 imaging sessions (part of the ICMS sessions; 4 and 3 imaging sessions from monkey L and A respectively). The comparison between visual stimulation and ICMS was done on 8 visual stimulation sessions and 8 ICMS sessions from monkey L (part of the ICMS sessions). Matlab software was used for all statistical analyses and calculations.

#### Basic VSDI analysis

The basic analysis of the VSD signal (Δf/f) was detailed elsewhere (Ayzenshtat et al. 2010; Slovin et al. 2002). Briefly, it consisted of choosing pixels with minimal fluorescence level (the threshold was set to 15% of maximal fluorescence enabled by the camera detector), then normalizing each pixel to its baseline fluorescence level (known as frame-zero division, Shoham et al., 1999) and finally, subtracting the average fixation-alone condition (i.e. blank condition) to remove the heartbeat artifact. This basic analysis removes in an unbiased manner most of the slow fluctuations originating from heartbeat artifact or dye bleaching within a trial (Shoham et al. 1999; Slovin et al. 2002). These steps are schematically illustrated and explained in Ayzenshtat et al. 2010 Fig. S12. In addition, we applied a 2D high-pass filter on the VSD maps to detect pixels located on/near blood vessels or on the stimulating electrode and remove them from analysis. Some of the VSD maps in the figures were low-pass filtered with a 2D Gaussian filter (typically we used σ=1.5-2 pixels) mostly for visualization purposes. Any filtration done during analyses is indicated specifically, in the Results text or figure legend.

#### Time course analysis of the VSD signal and size of activated area

In each recording session, the VSD maps were first averaged across trials for each condition and then divided by the blank condition (fixation alone; as explained above). The time course analysis and its characteristics were quantified for each recording session, separately. To detect the peak response amplitude in time and space we used the following steps: the VSD maps were low pass filtered on the spatial domain and the VSD signal of each pixel was then filtered on the temporal domain (sliding window of 50 ms). Next, we detected the peak response amplitude in time and space within a time window of 20-200 ms post stimulation onset, this was done for each visual area (V1, V2, V4). To compute the time-course of the evoked VSD signal for each area we defined a region of interest (ROI) for each area which included all pixels crossing a threshold of >=75% of peak response activation (e.g. Fig. 2–3). Time courses for ROI were computed by averaging the VSD signal across all pixels within the ROIs. Finally, quantitative analysis was done only on pixels with ICMS evoked response crossing an SNR threshold (2.5 STDs) from their baseline activity in each visual area (V1, V2, V4). The time to peak response (TTP) of the VSD signal for the ICMS condition was calculated for each ROI. Response latency of the VSD evoked response was defined as the first time point in which the VSD signal in the ROI crossed a threshold of 3 STDs from baseline activity, and remained above this threshold for the next 30ms. The distribution of VSD peak response (averaged within ±20ms around peak time response) in all ROI pixels was compared to that of the baseline activity. Finally the size of the cortical area activated by ICMS was quantified by the number pixels crossing a threshold of 3 STDs at peak response from their baseline activity multiplied by the pixel area.

**Figure 2:**
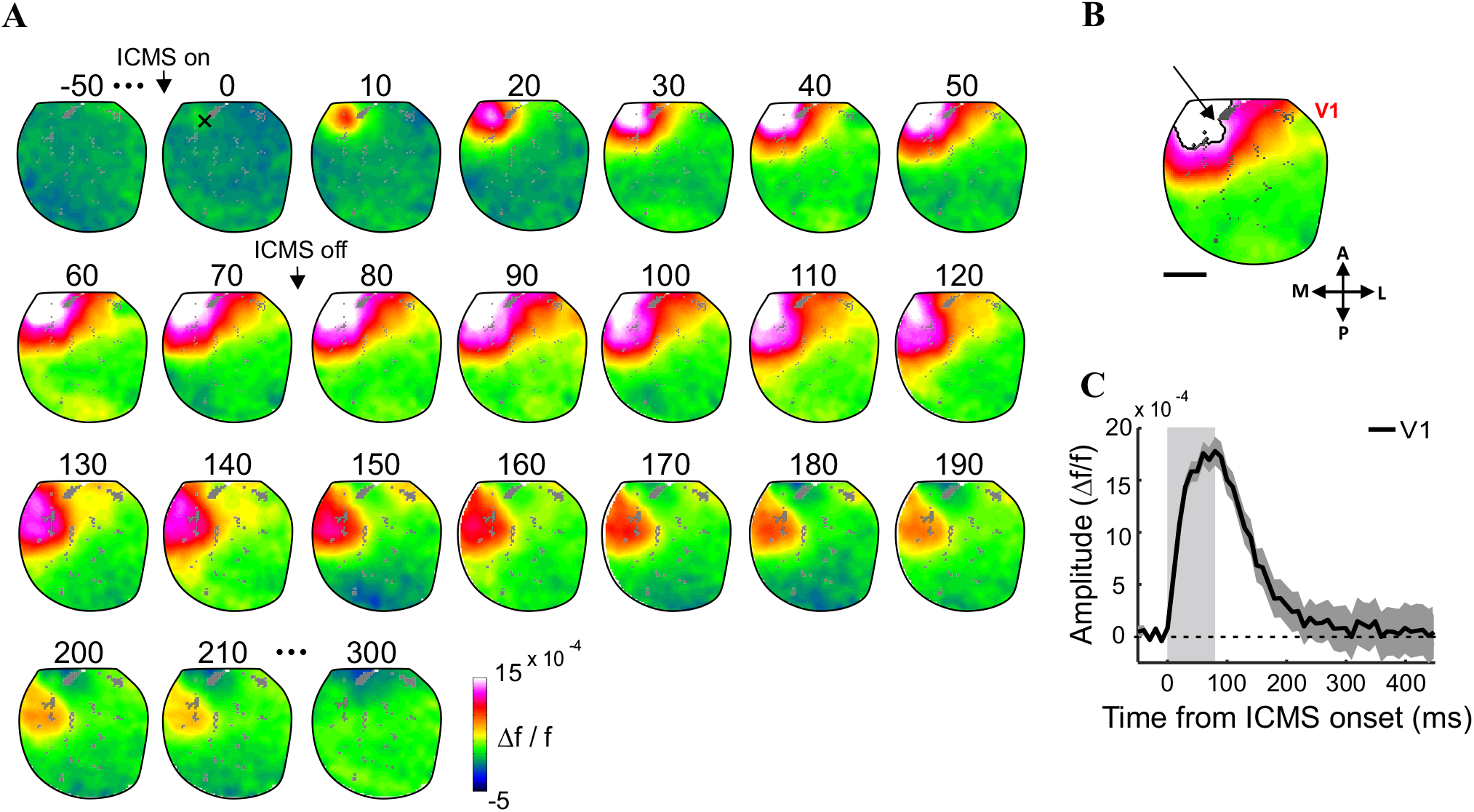
ICMS evoke population response in V1 that spread over few mm. **A.** A sequence of VSD i.e. population response maps from an example ICMS session (monkey L; TD 80ms), averaged over 26 stimulation trials (no stimulation). VSD maps were aligned on ICMS onset and filtered with a 2D Gaussian filter (σ=2pixels). The electrode position is marked with an X, black arrows denote ICMS on and off. **B.** Mean VSD map averaged over 60-90 ms after ICMS onset. The blue contour shows a Region of Interest (ROI) in V1 including pixels with response amplitude higher >= 75% of peak response in the recording session. The black arrow shows the microelectrode position. Gray pixels denote blood vessels (removed from analysis). Scale bar is 2 mm. **C**. Time course of the VSD signal averaged across the ROI pixels in (B). The gray rectangle area denotes microstimulation duration. Error bars denote ±1 SEM over trials.

**Figure 3:**
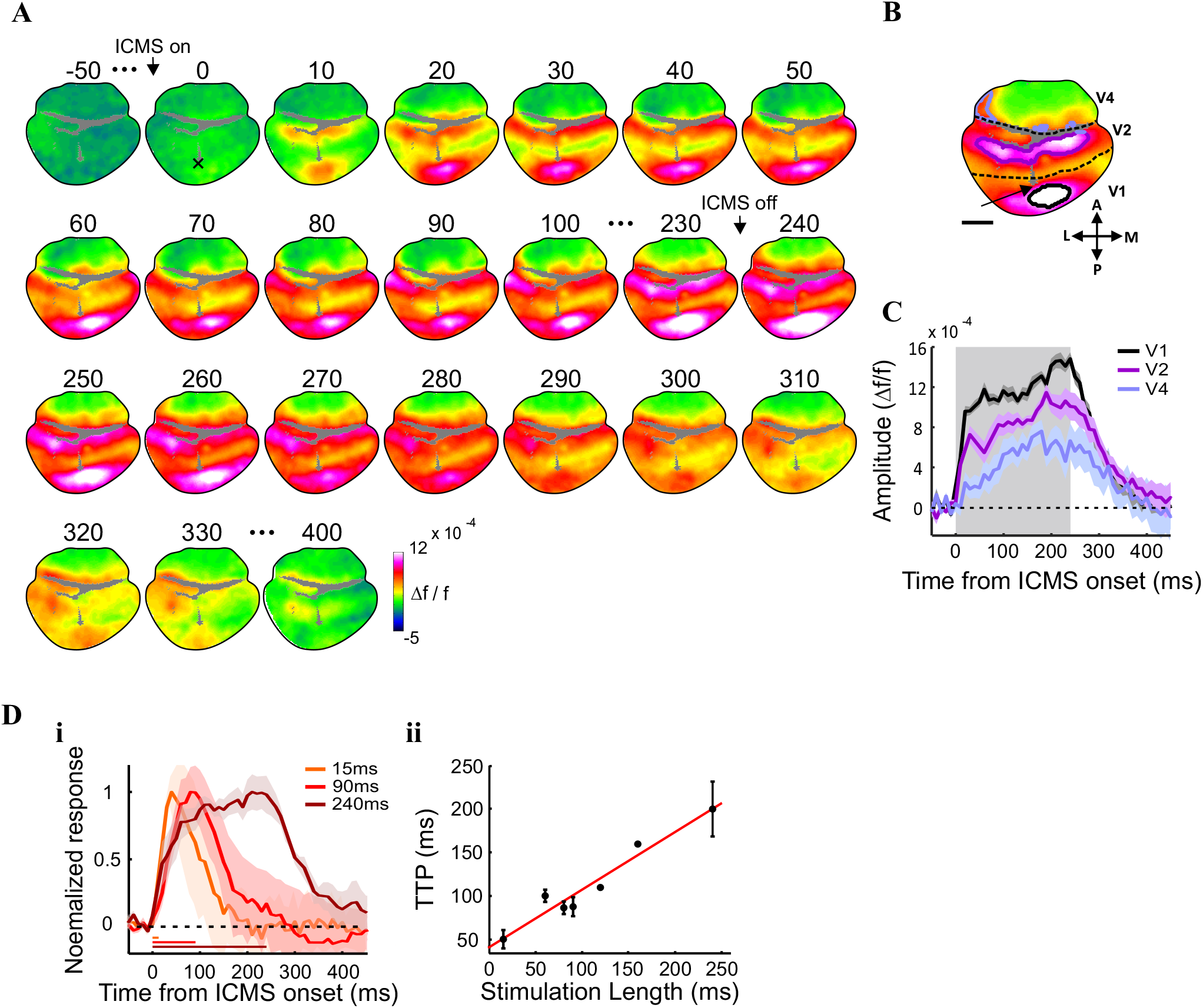
ICMS evoked population response can propagate from V1 to extra-striate cortex. **A.** A sequence of population response maps from an example ICMS session (monkey A; TD 240ms), average on 31 stimulation trials. VSD maps were aligned to ICMS onset. The electrode position is marked with an X, black arrows denote ICMS on and off. **B.** VSD map averaged over 210-240ms after ICMS onset with ROIs contours outlined on each area. The ROIs include pixels with response amplitude >=75% of peak response in V1 (black), V2 (purple) and V4 (light blue). Gray pixels denote blood vessels (removed from analysis). The border between V1 and V2 is denoted by a black dashed line. The lunate sulcus is denoted by the black dashed line between V2 and V4 area. **C.** Time course of the VSD signal averaged across the ROI pixels in (B). Gray area denotes microstimulation duration. Error bars denote ±1 SEM over trials. **Di.** Normalized time course for 3 representative TDs, averaged over imaging sessions. Time course were normalized to peak response. Shaded areas denote ±1sem over imaging sessions (TD=15ms, n=4; TD=90ms, n=4; TD=240ms, n=3). Horizontal colored bars denote stimulation duration. **Dii.** Time to peak (TTP) response as a function of stimulation duration, grand analysis. Data were averaged over sessions (TD=15ms, n=4; TD=60ms, n=3; TD=80ms, n=8; TD=90ms, n=4; TD=120 ms, n=1; TD=160ms, n=1; TD=240ms, n=3). Error bars denote ±1 sem. Red line shows a linear fit for the data points (r=0.97, p=0.0003).

#### Comparison between ICMS and visually evoked response dynamics

To compare the responses evoked by the two stimulation types, we studied the average VSD signal in the chosen ROIs (that include all pixels crossing a threshold of 75% of peak response activation in the recording session) from 8 ICMS sessions and 8 visual stimulation sessions during the rising and falling phases of the evoked responses. For the rising phase, the VSD signals were aligned on response onset and displayed only for the first 80ms, which was the stimulation duration in the ICMS sessions used for this analysis. TCs were normalized to peak response (of the ICMS or visual stimuli condition) up to 210 ms following stimulation onset. To quantify the falling phase, the VSD signal was aligned on the descending phase (defined as the time point with response amplitude lower by 3STDs from the peak response (ICMS) or steady state response (visual stim) for the next 30 ms and the VSD signal was normalized to the activity at this time point. Quantitative analysis was done only on pixels with ICMS or visually evoked response crossing an SNR threshold (2.5 STDs) from their baseline activity.

#### Comparison between ICMS and visually evoked response: Derivative maps and spatial profile

To compute the derivative maps we used the following equation (1):

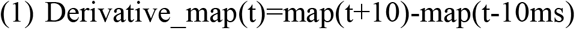

Where t is time in ms, and each frame duration is 10 ms long.

The derivative maps were first filtered using a low-pass 2D-Gaussian filter (Fig. 5A). Next we applied two orthogonal spatial profiles (width 10 pixels) crossing through the peak response (located in space; for the ICMS or visually evoked responses). The orientations of the spatial profile was set to be horizontal and vertical in the ICMS sessions. In the visual stimulation sessions, the spatial profile was set along the long axis of a fitted ellipse (to the 70% of peak response contour ROI) and orthogonal to that. The curves of the spatial profile (Fig. 5B) were computed by averaging along the width (10 pixels) of the profiles. Finally, to quantify the relation between the response amplitude at the center versus surround, we defined a measure for a center-surround modulation (CS-m):

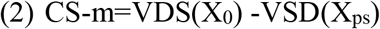

This was calculated on the curves of the spatial profile for the ICMS and visually evoked sessions, separately. CS-m was averaged across derivative time window corresponding to t=30-40 (Fig. 5B). X_0_ is the electrode position or to the spatial center of the visually evoked response in V1. The VSD values for X_0_ were averaged over 5 pixels, for the visual or ICMS responses. X_ps_ reflect the average VSD values in the proximal surround of the electrode, corresponding to ~1-2 mm away from X_0_ (range of 9-12 pixels away from X_0_).

#### VSD analysis of ICMS trials with evoked saccade

We first evaluated the site of the microstimulating electrode within the retinotopic map of V1, which was obtained in a separate set of experiments for each animal. For this purpose we used both optical imaging of intrinsic signal and VSDI while presenting the animal with small visual stimuli (7 and 6 sessions for Monkey L and A respectively). In 7 out of 25 ICMS sessions we found that the monkeys performed saccades directed to the retinotopic location of the electrically stimulated region in V1, and thus these saccades landed within the RFs of the stimulated neurons. Next we divided the ICMS trials into two categories: Class I. Trials where the animal performed a saccade that landed within the RF of the stimulated region in V1 (within a time window of 50-400ms post ICMS onset). For this class of trials we calculated the saccades’ amplitude, direction and latency (latency was defined as the time from ICMS onset until saccade initiation). Class II. Trials where the animal did not make saccades to the stimulated RF. Next, we computed the VSD response within a ROI located around the electrode for trials with saccades to the RF (class I) and compared them to trials without saccades to RF (class II). The VSD response was aligned on ICMS onset and data was normalized in each trial by subtracting the baseline activity and then dividing with the peak response of every ROI. The ROI included pixels that crossed a threshold of at least 70% from the maximum amplitude difference between class I and class II trials.

### Statistical analyses

Nonparametric statistical tests were used: the Wilcoxon rank-sum test to compare between two medians from two populations or the Wilcoxon signed-rank test to either compare a population’s median to zero or compare the median of differences between paired samples to zero. Data in the Results section are presented in mean±sem or as median±mad unless specified otherwise.

## Results

We asked what are the spatio-temporal population responses patterns evoked by intra-cortical microstimulation (ICMS) in the visual cortex of alerts monkeys and what is the relation of these activation patterns to those evoked by visual stimulation. To investigate this, we combined voltage-sensitive dye imaging (VSDI) in fixating monkeys with ICMS or visual stimulation (Fig 1A-B). A microelectrode was inserted in the upper layers of V1 area (Fig1B, left) and short trains of ICMS were applied while the animals were maintaining fixation on a small fixation point. The ICMS parameters we used (Fig. 1B middle; see Methods) were previously shown to be highly effective in evoking neural activity in visual cortices as well as in generation of saccades directed to the RF of the stimulated neurons and affect behavioral responses or visual perception (Bradley et al. 2005; Davis et al. 2012; Schiller et al. 2011). We then used VSDI to measure and characterize the ICMS or visually (Fig. 1B right - for visual stimulus) evoked population responses at high spatial (meso-scale; 100×100 μm^2^/pixel) and temporal resolution (100 Hz). The fluorescence dye signal (Δf/f) from each pixel reflects the sum of membrane potential changes of all neuronal elements (dendrites, axons, and somata) and therefore reflect population activity rather than responses of single neurons (Slovin et al. 2002). In addition, the VSD signal emphasizes subthreshold synaptic potentials (but reflects also suprathreshold activity; Ayzenshtat et al., 2010; Jancke, Chavane, Naaman, & Grinvald, 2004). A total of 33 VSDI sessions were analyzed from two hemispheres of two adult monkeys (see Methods).

### ICMS evoked neural activity that spread horizontally within V1 and could propagate to extrastriate cortex

Figure 2A shows data from an example session: a sequence of VSD maps evoked by a short ICMS train duration (TD, 80 ms). The population response in V1 appeared first around the electrode site with a very short latency of 10 ms after stimulation onset. In subsequent time frames (each frame is 10 ms) the population activity quickly spread horizontally within V1. The VSD response arrived to peak response amplitude at the end of the train duration. The size of the activated area at peak response, was quantified from the number of activated pixels crossing a threshold (see Methods) and was 4.3^2^ mm^2^. At the end of stimulation, the population response decreased back to baseline activity within the next 200 ms. Fig. 2B shows the VSD map averaged around peak response activation: a clear patch of increased population activity is evident near the electrode site. The blue contour depicts a Region of Interest (ROI) including all V1 pixels with response amplitude equal or larger than 75% of peak response (see Methods). Fig. 2C shows the time course of the VSD signal averaged across the ROI’s pixels: following stimulation onset, the population response shows a fast rising phase that is followed by second short phase which appears to increase at a slower rate. The population response peaked at the end of train duration (80-90 ms; 1.7×10^−3^ Δf/f, p=6.55×10^−10^ for difference from baseline activity, Wilcoxon rank-sum) and then descended back to baseline at a much slower rate relative to the rising phase.

Figure 3A shows an example session from a different animal, where we were able to image several visual cortices simultaneously: V1, V2 and V4. The VSD response started to increase within 10 ms after stimulation onset and appeared first around the electrode site in V1 and also in V2 area. In subsequent time frames the population activity spread horizontally within V1 and V2 areas and started increasing in area V4 with a small visible delay relative to V1 and V2. The VSD response arrived to peak response amplitude in all areas around the end of the train duration (240 ms) with the highest activity in V1 and lower response amplitude in V2 and V4. The activated region was of similar size: ~4^2^ mm^2^ in V1 and V2 with a smaller area of 2.5^2^ mm^2^ in V4 (note that part of V2 is in the lunate sulcus and not available for imaging). At the end of stimulation, the response decreased to baseline activity within the next 200 ms. Fig. 3B shows the mean VSD map averaged peak response (210-240 ms after ICMS onset). Clear separated patches of activation are shown within each of the visual areas denoted by the colored ROI contours. Interestingly, the distinct patches of activation in V1 and V2 appear to be spatially separated despite their spatial proximity and short distance from the stimulating microelectrode and the vertical meridian (the latter is also the border between V1 and V2; Fig. 3B dashed black line). This result may suggest that the ICMS evoked neuronal activity that spread mainly through functional connections in V1 and V2, i.e. local horizontal connections within each area as well as feedforward connections between V1 and V2. This may also suggest that the activation patches in V1 and V2 reflect retinotopically matched regions i.e. similar location in the visual field. In other words, it is possible that the neural activity in V1 propagate to V2 through feedforward projections from V1. The colored contours in Fig. 3B depict the ROIs around peak response in V1, V2 and V4 respectively. Fig. 3C shows the time courses of the VSD signal for each ROI: following stimulation onset, population response shows a fast rising phase followed by a much slower increasing phase. The population activation peaked around the end of train duration (240 ms), with the highest activity in V1 (1.4×10^−3^ Δf/f), lower amplitude in V2 (1.0×10^−3^ Δf/f) and V4 with the lowest response amplitude (6.8×10^−4^ Δf/f). Following end of stimulation, the VSD response decreased back to baseline at a rate that was much slower than the rising phase. The examples in Figures 2 and 3 suggest that ICMS in V1 evoked neural activity spreading horizontally within V1 surface over several mm and that activity can also propagate to extrastriate cortices (V2 and V4).

The grand analysis across all recording sessions confirmed the results depicted in the two example sessions and showed that the median latency for response onset in V1 was 10 ms (±10, mad; n=25) and the neural activation at peak response, covered a mean area of 4.2^2^ mm^2^ (±0.3^2^ sem; n=25; significant VSD response was observed in all ICMS sessions). In Area V2, the median latency to response onset was 10 (±0, mad; n=5) which was not significantly different from the median latency in V1 (p=0.3; Wilcoxon rank-sum; however we note that the temporal resolution we used is 10 ms). The mean size of activated region in V2 was 3.9^2^ mm^2^ (±0.27^2^, sem; n=5; not significantly different from V1; p=0.74; Wilcoxon rank-sum). The median response latency in area V4 was 35 (±10, mad; n=4; significantly longer than V1; p=0.016; Wilcoxon rank-sum) covering a mean area of 2.0^2^ mm^2^ (±0.6^2^, sem; significantly smaller than V1; p=0.01; Wilcoxon rank-sum). Figure 3Di shows the time course of the VSD signal (averaged across imaging sessions) from three representative train durations (15, 90 and 240 ms). Following ICMS onset the VSD response showed a fast rising phase of activation, and for longer train durations (90, 240 ms) this was followed by a slower increasing phase that lasted until end of stimulation. Finally, there was a strong linear correlation between the time to peak response (TTP) of neural activation in V1 and the train duration within the used range (15-240 ms; Fig. 3Dii; r=0.97, p=0.0027). Notably, the TTP for very short trains (15 ms) was approximately 3 times longer (50 ms) than the train duration itself, however for longer train duration, the TTP was more similar to the train duration. In summary the results above suggest that when electrically stimulating V1, the response spread horizontally across several mm in V1 and can propagate to extra-striate areas.

### Comparison between the visually and ICMS evoked activity in V1

Electrical stimulation in human’s V1 was previously reported to elicit the visual sensation of a small flickering spot of light termed as phosphene. The position of the phosphene in the visual field corresponds to the stimulated region in the retinotopic map i.e. the location of the stimulated RFs in the visual field. Previous studies suggested that similar phenomena occur when stimulating V1 of behaving monkeys (Tehovnik and Slocum 2007). However, the degree of similarity between the neural activation patterns evoked by ICMS and those evoked by naturally by small visual stimuli remains elusive. To investigate this, we compared the spatio-temporal patterns of population responses evoked by a small visual stimulus to those evoked by ICMS. Figure 4A shows the VSD maps evoked by a small Gabor patch while the animal was performing a fixation task (see Methods). As expected, due to the propagation speed of neural activity from the retina to V1, VSD signal started to increase ~50-60 ms after visual stimulus onset. The median latency of population response in V1 across all visual sessions (n=8) was 55 ms (±5, mad) which was significantly longer than the response latency for ICMS (p=0.009; Wicoxon rank-sum, 8 ICMS sessions). Figure 4A shows that the visually evoked response showed increased local activation combined with horizontal spread of activation over few mm in V1, similar to the ICMS evoked responses. As for the ICMS condition, the VSD map at peak response amplitude (Fig. 4B) shows a clear patch of activation and the VSD time course (Fig. 4BC) shows a rising phase that is faster than the descending phase. The grand analyses across all visual stimulation sessions shows that the mean activated area at peak response evoked by the small Gabor was 6.1^2^ mm^2^ (±0.3^2^, sem; n=8). This was significantly larger than the mean area activated by the ICMS 4.2^2^ mm^2^ (±0.3^2^, sem; p=0.035 Wicoxon rank-sum). This result may suggest that the size of the generated phosphene was smaller than the size of a 0.5° high contrast Gabor (see Discussion).

**Figure 4:**
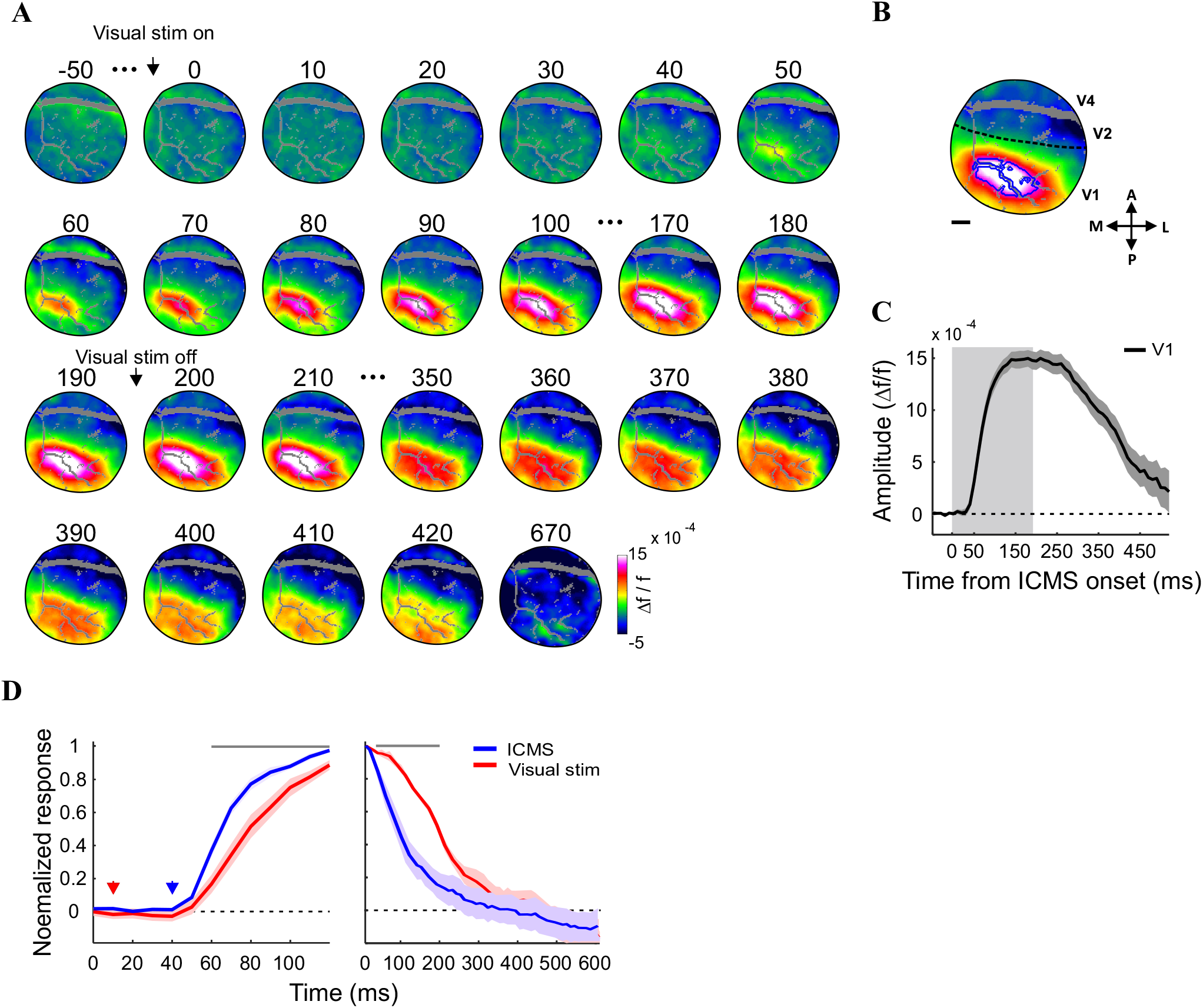
Visually evoked responses and the comparison of their time course dynamics to ICMS evoked responses. **A.** Population response maps of an example visual stimulation session averaged across12 trials (monkey L). The VSD maps are aligned on visual stimulus onset (high contrast Gabor patch, 2σ=0.25; stimulus duration: 200 ms). **B.** VSD map averaged over 180-230ms after visual stimulation onset. Blue ROI denote V1 pixels with response amplitude higher than 75% of peak response in the recording session. Gray pixels denote blood vessels (removed from analysis). **C**. VSD time course of the response in the ROI. Gray area denotes visual stimulus duration (200 ms). Error bars denote ±1 sem over trials. **D.** Left: Comparison of the initial rising phase of the VSD signal evoked by ICMS (red curve, TD=80ms, n=8 sessions) and by visual stimulation (blue curve, n=8 sessions). Time courses were aligned on response onset for both stimulation conditions and are plotted and compared for the first 80 ms (to compare similar stimulation length of ICMS and visual stimulation). VSD time courses were normalized to peak response of each session. Red and blue arrows denote visual stimulation onset and ICMS onset, respectively. Right: Same as in Left, but responses were aligned on turning off the ICMS or turning off the visual stimulation and normalized to that point in time. Error bars denote ±1 sem over sessions.

To compare the time course dynamics of the visually and ICMS evoked responses we compared the rising and the descending phases of the evoked neural activation for both stimulation types. We aligned the VSD signal on the latency onset of the VSD rising phase for both ICMS and visual stimulation or on the descending phase (see Methods). Fig 4D left shows the comparison of the rising phase aligned on response onset (plotted for the first 80 ms for both stimulation conditions). The VSD signal evoked by ICMS shows a faster rising phase than that evoked by visual stimulation and the top gray bars indicate the time points where the ICMS showed a significantly higher amplitude than the visually evoked response (p-values varied between p<0.05 to p<0.001 for the different time points; Wilcoxon rank-sum). Figure 4D right shows the VSD signal aligned on the initiation of the falling phase and in this case the VSD signal also decays faster for the ICMS condition. The top gray bar indicates all time points where the ICMS showed a significantly lower value (p-values varied between p<0.05 to p<0.001 for the different time points; Wilcoxon rank-sum) than the visually evoked response. Thus for both the rising and falling phase of the VSD evoked response, the ICMS condition showed faster dynamics (similar results were obtained for ICMS of similar time duration to the visual stimulation).

The faster dynamics of the ICMS evoked responses as compared with the visually evoked responses suggests that the spatio-temporal activation patterns of the two stimulation conditions may show some differences. To investigate this, we computed the VSD derivative maps (from each VSD map we subtracted the VSD map measured 20 ms earlier, see Methods) for the visual and ICMS conditions and then aligned the derivative maps on the population response onset. Example derivative maps for the two conditions are depicted on Fig. 5A. The first derivative map (t=0) in the visual stimulus condition (Fig. 5Ai) shows the *change* in the spatial pattern of the VSD signal when the evoked response started to develop, revealing a small local patch of increased population response. This patch is like to reflect the thalamic input arriving to V1, activating the neuronal population within the retinotopic map corresponding the location of the stimulus in the visual field. The following two derivative maps (t=10, 20) shows additional and larger positive change in the VSD signal: the local patches in the derivative maps show higher activation change which spreads horizontally over the cortical surface. This change is attenuated in the last three frames (t=30-50), which show a smaller VSD derivative signal as the evoked VSD response approaches a plateau (see Fig. 4C). Interestingly, the derivative maps for the ICMS condition reveal some differences in the spatial pattern. The first two maps (t=0,10) are similar to the visual stimulus condition and as expected, show the appearance of a local patch of increased population response near the electrode site, which seems to spread horizontally over the cortical surface. The next few derivative maps show a spatial pattern that is different from that of the visual stimulus condition. At t=20-50 ms (from response onset), the VSD derivative map reveals an annulus shape pattern around the electrode site, as evident by a red-orange ring with yellow-green hole in the middle. This means that the derivative signal at the electrode site shows lower values as compared to the proximal regions located ~1-2 mm away from the microelectrode, that have higher values. This results is different from the derivative maps of the visually evoked responses, demonstrated a more “homogeneous” or Gaussian like patch of activation change (Fig. 5Ai, t=30-50). These results were further quantified across all visual sessions (n=8) and ICMS sessions (n=8) by computing the spatial profiles (see Methods). Fig. 5B shows the grand average (across all sessions) of the spatial profiles, for the visual stimulus condition (Fig. 5Bi) and ICMS condition (Fig 5Bii; note that spatial profiles for both condition were truncated at the border of the imaged area). The spatial profiles in the ICMS condition show a positive hill of change (t=0, black curve) that is centered on the electrode site (distance=0, x-axis). At t=10 the derivative signal (brown curve), increases in amplitude and spatial spread. Interestingly, at t=20, 30 (dark orange and orange curves) the spatial profile of the derivative signal shows a more complex spatial pattern which reveal two hills that are separated by a trough, the latter positioned at the microelectrode site. Thus, the derivative signal at the electrode position (distance=0 on the x-axis) shows lower values relative to regions at the proximal surround (distance=1-2 mm away from the microelectrode). Moreover the derivative profile seems to further expand horizontally, over the cortical surface. Thus the derivative maps analysis suggests that in the ICMS condition the VSD signal at the electrode site arrived to peak or plateau activation earlier than regions in the proximal surround which seems to lag behind and show a slower horizontal propagation of neural activation. In contrast, the derivative signal for the visually evoked condition, shows a rather homogeneous, gaussian like hill of activation (Fig. 5Bi). To further quantify this effect, we plotted in Fig. 5C the VSD derivative value at the electrode location (distance=0; x-axis) vs. the VSD derivative value at 1-2 mm distance from the electrode location (y-axis; Figure 5C). In the ICMS condition, the VSD derivative values are mostly above the diagonal line (denoting identical amplitude values for the center and surround). This means that for most ICMS recording sessions, the VSD derivative was lower at the electrode site vs. the proximal surround (1-2 mm away). Next, we defined a center-surround modulation measure (CS-m; see Methods) as the difference between the VSD derivative at center (i.e. microelectrode location or center of visually evoked response) versus the mean derivative value at x=1-2 mm away from the center. The mean CS-m for the ICMS was negative −1.3×10^−4^ (±2.2×10^−5^, sem; n=16 spatial cuts) and significantly smaller from zero; p=0.00053, Wilcoxon signed-rank). In contrast, in the visual condition, the derivative values are mostly below the diagonal (Fig. 5C), meaning that for most recording sessions the VSD derivative at the x=0 was higher than in the proximal surround regions, 1-2 mm away. The mean CS-m was positive for the visual condition 9.4×10^−5^ (±1.9×10^−5^, sem; n=16 spatial cutes) which was significantly larger from zero (p=0.0064). Finally, Fig. 5C shows a clear and significant separation between the ICMS values (blue symbols) and visually evoked values (red symbols). The mean distance from the diagonal for the ICMS was 8.9×10^−5^ (±1.6×10^−5^, sem) which is significantly different (p=2.25×10^−6^; Wilcoxon rank-sum) from the mean distance for the visual evoked response: −6.7×10^−5^ (±1.3×10^−5^, sem). These finding may suggest the involvement of an inhibitory process at the electrode site in the ICMS condition relative to the visually evoked condition.

**Figure 5:**
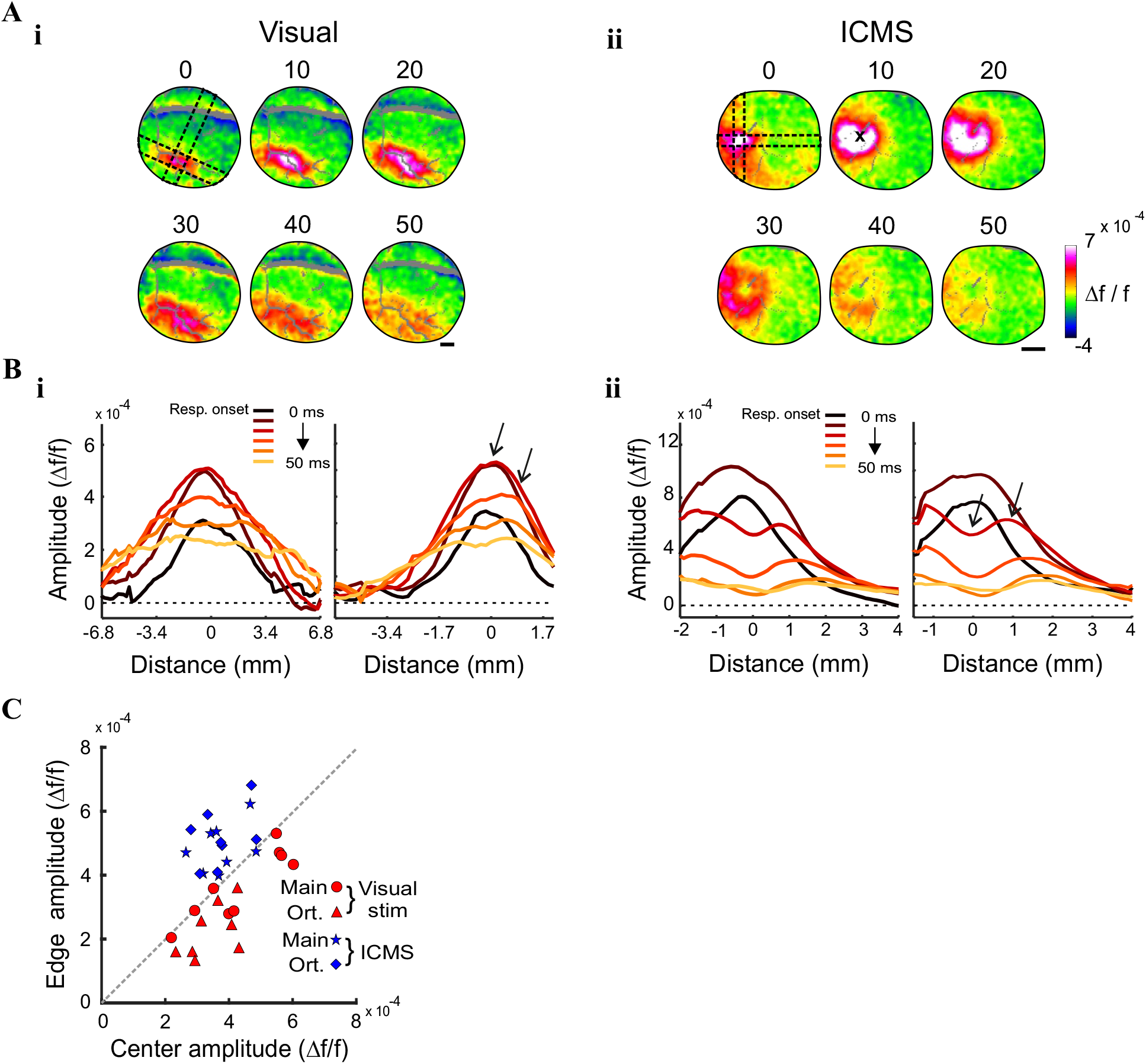
Derivative maps and spatio-temporal changes in spread of visually and ICMS evoked responses. **A.** A sequence of derivative VSDI maps, example sessions. i: Derivative maps for visually evoked responses (same session as Fig. 4); ii: Derivative maps for ICMS evoked response (average over n=34 stimulated trials; TD=80ms; monkey L). Each map represents the VSD difference of two subsequent frames (see Methods). Time above each map is denote time from response onset (see Methods). Maps were filtered with a 2D Gaussian filter (σ=2 pixels). Black dashed contours show the location of the spatial profiles for the example sessions. Gray pixels denote blood vessels (removed from analysis). X denote microelectrode position. **B.** Grand average (across all sessions) of the derivative spatial profiles (depicted by dashed black lines in A; see Methods), at increasing time from response onset. i: visually evoked responses (n=8 recording sessions). left: spatial profiles in the main axis of activation, right: spatial profiles in the orthogonal axis. ii: same as in i, but for ICMS evoked responses (n=8 recording sessions, TD=80). Note that the spatial profile are not symmetric because they were truncated at the edges of the imaging chamber. **C.** The VSD response amplitude at the center of the spatial profile (x-axis) vs. the VSD response amplitude at proximal regions, 1-2 mm away from center (y-axis). ICMS sessions are denoted with blue marker (n=16 for main and orthogonal spatial profiles) and visual sessions are denoted by red markers (n=16 for main and orthogonal spatial profiles). Black arrows in B depict approximately the locations of the center and proximal surround regions.

### ICMS evoked saccades and the relation to evoked spatio-temporal maps

Next, we were interested to investigate the ICMS trials in which the animal made saccades to the stimulated RFs. In 29% of the ICMS sessions (7/25 sessions; see Methods) we found that following stimulation onset, the animals made saccades towards the location of the stimulated RFs (the retinotopic maps of the imaged area were obtained in a separate set of experiments, see Methods). Figure 6 shows two example sessions of saccade trajectories and their characteristics from two animals. Figure 6A,B shows the trajectory of the evoked saccades in the stimulated trials (left panels) while the non-stimulated trials (right panels) show no evoked saccades towards the stimulated RFs. The distribution of saccade size for the examples in Fig. 6A,B are depicted in Fig 6C,D left panels. The mean size of the evoked saccades was 1.44° (±0.08, sem; n= 24 trials) and 1.62° (±0.06, sem; n= 33 trials) for monkey L and A respectively. The mean saccade latency was 202 ms (±22, sem) and 220 ms (±15, sem) in the two examples (Fig. 6C, D right panels). The grand analysis across all sessions (n=7), shows that the mean saccade size and mean latency was 1.5° (±0.05, sem) and 219 ms (±9, sem). Additional analysis revealed that within the above sessions (n=7) only a fraction of the stimulated trials (62% for both animals) have shown evoked in saccades to the stimulated RF (57/81 trials in monkey A and 75/139 stimulated trials in monkey L). We then hypothesized that the population activity in trials w/saccades was larger than the population responses in trials w/o saccades. We therefore decided to investigate the VSD signal in stimulated trials with evoked saccades versus stimulated trials w/o evoked saccades. Figure 7A shows an example session: a sequence of VSD maps averaged across stimulated trials with evoked saccades. Following ICMS onset (t=0), a clear patch of increased population response appears near the electrode site and activation then spreads over few mm within V1. Figure 7B shows the same example session, however, here the VSD maps were averaged across stimulated trials that *did not* show evoked saccades to the stimulated RF. The VSD response in Fig. 7B shows a population response that seems to expand over a smaller area and shows a lower response amplitude in most of V1 area. To further quantify this we computed the VSD signal for the data in Fig. 7A-B in a ROI near the electrode site and aligned it on ICMS onset. Figure 7C shows the VSD time course for stimulated trials with saccades (black curve) and trials w/o saccades (grey curve): interestingly the VSD signal shows a higher population response for trials with saccades (p=0.02; Wilcoxon rank-sum test over trials). Next, we applied this analysis separately for each animal (Fig. 7D) and the grand analysis confirm the example shown in Fig. 7C. Figure 7D shows that the population response was significantly higher for trials with saccades vs. trials w/o saccades for both animals. For monkey L (Fig. 7D left) the mean normalized response in a time window of 30-130 ms following ICMS onset for trials with evoked saccades was 0.82 (±0.05, sem; n=27 trials) compared with 0.57 (±0.05, sem; in trials w/o saccade, n=21 trials; p=0.003, Wilcokson rank-sum test). For monkey A, the mean normalized response in a time window of 40-140 ms following ICMS onset for trials with evoked saccades was 0.77 (±0.04, sem; n=73 trials) compared with 0.577 (±0.05, sem) in trials w/o saccades (n=56 trials; p=0.00008; Wilcokson rank-sum test). In summary, the population response in trials with saccades showed a response amplitude that was higher by 25-30% as compared with trials with w/o saccades (note that the saccades themselves initiated at much later time).

**Figure 6:**
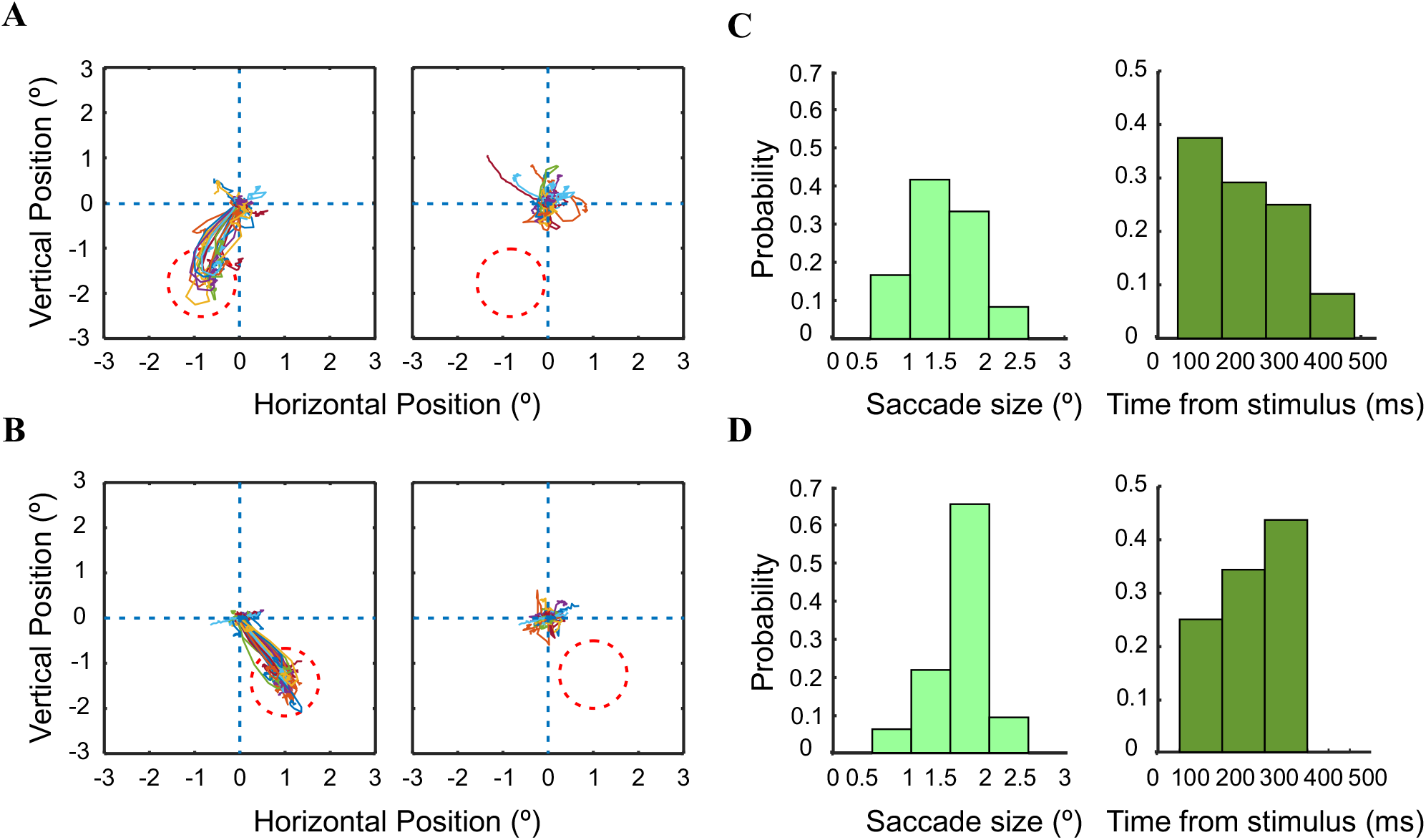
ICMS evoked saccades. **A.** Eye position trajectories for an ICMS example session, monkey L. Left: eye position trajectories in stimulated trials with a saccade targeted to the stimulated region in the retinotopic map (red dashed circle; n=24 trials). Right: eye position trajectories in non-stimulated trials (n=32 blank trials). **C.** Histograms of saccades size (left) and saccade latency from microstimulation onset (right). **B&D.** Same as A&C but for an ICMS example session from monkey A. (n=33 trials with saccade and n=36 blank trials).

**Figure 7:**
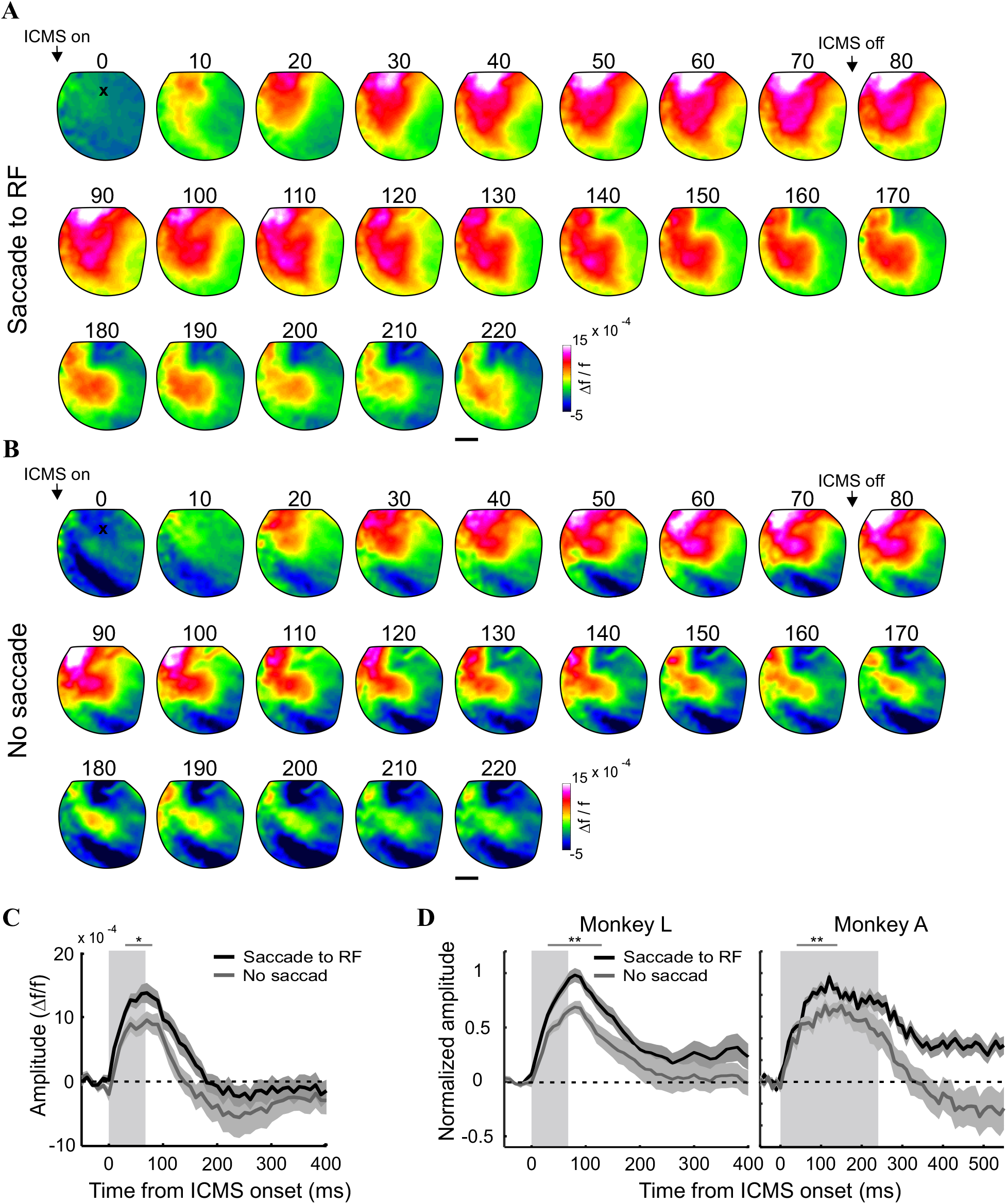
Trials with saccades directed to the stimulated RF demonstrate a higher VSD signal. **A&B.** Sequence of VSD response maps for an example session (same as Fig. 6A; TD=80ms). VSD maps were aligned on ICMS onset and filtered with a 2D Gaussian filter (σ=2 pixels). **A.** VSD maps for trials with saccades directed to the stimulated site region in V1, i.e. to the stimulated RFs in V1. Maps were averaged over 24 trials. **B.** Same imaging session as in A, but for trials w/o saccades. VSD maps averaged over n=8 trials. **C.** Time course of the VSD signal in a ROI near the microelectrode site. Black and grey curves denote trials with saccades and w/o saccades, correspondingly. Rectangle gray area denotes stimulation duration. Error bars denote ±1 SEM over trials. The top gray horizontal line denotes the time window used to compute statistical significance between the two VSD time courses (*p=0.02 Wicoxon rank-sum test). **D.** Left: time course of the VSD signal in monkey L, averaged over trials with saccades (n=57; black curve) and trials without saccades (n=21 trials; gray curve). Rectangle gray area denotes microstimulation duration. Error bars denote ± sem over trials. The top gray horizontal line denotes the time window used to compute statistical significance between the two VSD time courses (**p=0.0019; Wicoxon rank-sum test). Right: same as left but for monkey A. n=73 trials with saccades and n=56 trials without saccades (***p=0.00008).

## Discussion

One of the major goals of microstimulation in V1 is to study the feasibility of cortical visual neuroprosthesis and the development of useful artificial vision at high resolution for blind human patients (Lewis et al. 2015; Lowery et al. 2015; Tehovnik et al. 2009). For this goal to be realized, one needs to characterize the neural and behavioral responses evoked by ICMS and compare it with neural responses evoked by visual stimulation. Using VSDI in fixating monkeys we studied the spatio-temporal patterns of neural activity evoked by ICMS in V1, at high spatial (meso-scale) and temporal resolution. We characterized the evoked neural activation and compared it to population responses evoked by visual stimulation. Finally, we also investigated the neural activity when the animals made saccades to the stimulated RFs, following microstimultion onset, in comparison to trials w/o evoked saccades.

### Spatio-temporal properties of V1 response to ICMS

Although several groups investigated the effects of microstimulation in V1 of behaving monkeys, most of the reported results were at the behavioral level (Bartlett and Doty 1980; Bradley et al. 2005; Davis et al. 2012; Doty 1965; Tehovnik et al. 2003). Tolias et al. (2005) combined fMRI with microstimulation in anesthetized monkeys to estimate the spread of neural activation in V1 and its propagation. However, the BOLD signal does not directly measure neural activity and the hemodynamic response is much slower (second resolution) as compared with neuronal scale (millisecond resolution). Thus, little is known on the spatio-temporal patterns of neuronal responses evoked by ICMS in behaving monkeys. Using VSDI we were able to measure and characterize the spatio-temporal patterns evoked by ICMS onset, at high spatial (meso-scale) and temporal resolution.

Within the parameters used in our work, ICMS induced widespread cortical activity that spread over an average area of 4.2^2^ mm^2^ in V1 at peak neural response. This value is a much larger from expected passive current spread that can cause direct excitation of cortical elements (Tehovnik et al. 2004). The large spread of activation in V1 reported in our study is in accordance with previous reports in monkeys (Brock et al. 2013; Histed et al. 2009; Tolias et al. 2005). This result suggests that the activated area at peak neural response emerged not only from direct activation of the current spread but reflects the spread of neural activation through horizontal connections and local synaptic interactions (Fehérvári et al., 2015). Additional factors that can contribute to the large spatial spread could be the electrical excitation of axons passing near the stimulation site and also feedback effects arriving from V2 or V4 (Klink et al. 2017). The size of the activated area we measured in response to microstimulation is somewhat smaller than what was previously measured using fMRI (Tolias et al. 2005), however Tolias et al. used much higher currents and much longer stimulation durations.

Our results show that ICMS in V1 evoke local neuronal activity that can propagate to higher visual areas (V2, V4; Fig. 3), in accordance with previous studies in monkeys suggesting multi-synaptic propagation (Klink et al. 2017; Matsui et al. 2011). This is also in accordance with the reported influence of microstimulation on behavioral choices and perceptual decisions implying an impact on high-level decision stages (Gold and Shadlen 2007; Salzman et al. 1990; Uka and DeAngelis 2006). Although previous studies reported that ICMS in V1 or LGN can propagate downstream, to areas that are only one synapse away (Logothetis et al. 2010; Sultan et al. 2011; Tolias et al. 2005), the longer response latency in V4 (relative to V2) suggests that neural activity propagated to V4 via a multi-synaptic pathway (and not directly from V1 projections(Yukie and Iwai 1985)). Our results are also in accordance with previous ICMS studies in rodents which applied similar ICMS parameters in V1 and reported on local, wide spread population activity that propagated to higher visual areas (Fehérvári et al., 2015, 2016; Dadarlat et al. 2019)). As the temporal resolution we used our experiments is 10 ms, the latency of the measured response in extrastriate cortex cannot determine the exact propagation circuitry. The activation in V2 can be caused by corticocortical single synaptic connection or it can be caused by polysynaptic signal propagation (cortico-thalamo-cortical connections (Bender, 1983; C. Hilgetag, Burns, O’Neill, Scannell, & Young, 2000; C. Hilgetag, O’Neill, & Young, 2000; Rushmore, Payne, & Lomber, 2005). Moreover, due to the temporal resolution we used we cannot distinguish between orthodromic and antidromic activation of V2.

### Comparison between ICMS and visually evoked neural responses

It is well established that electrical stimulation of V1 induces perception of light sensation that resembles small visual stimuli (Brindley and Lewin 1968; Clark et al. 2011; Foerster 1929; Penfield and Perot 1963). However, the relation between the neural activity patterns evoked in V1 by visual stimulation and ICMS is not well understood. To investigate this, we compared the activity in V1 evoked by both stimulation types. While both small visual stimulus and ICMS induced a local patch of neural activation, there were several differences on both the temporal and spatial dimensions.

ICMS and small visual stimuli activated V1 region extending over tens of mm^2^ and in both cases we observed lateral propagation of neural activation, suggesting the involvement of horizontal connections (Fehérvári et al., 2015; Slovin et al., 2002). Interestingly, it was previously reported that ICMS stimulation in V1 of NHPs within the central visual field generates the perception of small phosphenes with an estimated size of 0.17-0.5° in diameter (Schiller et al. 2011). These values are in accordance with our neurophysiological results that the spatial spread of electrically evoked activity is within the range or smaller than the area activated by a visual stimulus size of 0.5°. Visual stimulation induce VSD activation in V1 which is then propagated to V2 and V4 (Ayzenshtat et al. 2010; Meirovithz et al. 2010; Slovin et al. 2002). Similarly, in the ICMS condition, we also observed propagation of neural activity into downstream areas to V1, i.e. V2 and V4 areas. We previously reported that a single pulse of ICMS in the barrel cortex of anesthetized rats has generated similar spatio-temporal response patterns to those evoked by a single whisker deflection (Nivinsky Margalit and Slovin 2018). ICMS in V1 of monkeys was shown to activate local functional networks or functional network along the visual pathway (Brock et al. 2013; Klein et al. 2016; Klink et al. 2017). Furthermore optostimulation in monkey’s V1 showed normalization attributes when given in conjunction with a visual stimulus, in a similar manner to that of two visual stimuli given together (Nassi et al. 2015). Lim et al. (2012) also reported very high similarities between optogenetically evoked and sensory evoked responses in several primary sensory areas in the mouse cortex, including the visual cortex. Finally, several human studies reported on the similarities between electrical stimulation and visual sensations (Beauchamp et al. 2018; Winawer and Parvizi 2016).

As expected, due to the propagation velocity of neural activity from the retina to V1, the ICMS response had a much shorter latency (~10 ms) compared with the visually evoked response (40-50 ms). These results are in accordance with previous reports in the visual cortex, both in rodents (Fehérvári & Yagi, 2016; Fehérvári et al., 2015) and NHPs (Klink et al. 2017). Additional analyses revealed that when aligning the VSD signal of both stimulation conditions on the neural response onset, the rising response and the falling response of the ICMS condition showed faster dynamics when compared with the visual stimulation. This is in accordance with previous publications in rodents (Butovas & Schwarz, 2003; Fehérvári & Yagi, 2016; Fehérvári et al., 2015). A possible explanation for this is that ICMS induced a highly synchronized pulse of neuronal activation, leading to a fast increase in the VSD signal, which is a population signal.

The VSD derivative maps analysis suggests that in the ICMS condition the VSD signal at the microelectrode site arrived to peak response faster than proximal regions around the microelectrode, which lagged behind. Moreover, the annulus shape pattern with a hole in the middle, in the derivative maps (Fig. 5A) suggested a travelling wave of activity, emerging at the electrode site and propagating horizontally through the proximal regions around the microelectrode. This spatial patterns may suggest the involvement of inhibitory processes such as GABAA receptors (Fehérvári et al., 2015) activated with a varying gain over the spatial domain. Additional previous publications reported on inhibitory activity evoked by ICMS (Butovas and Schwarz 2003; Dadarlat et al. 2019; Klink et al. 2017). The derivative maps of the visually evoked condition showed a more homogeneous pattern of activation spread, and lacked the pattern of an annulus with a hole shape. Finally, we did not check systematically over ICMS parameters and it reasonable to assume that the patterns overserved in the derivative maps are depended on the ICMS parameters.

### ICMS and evoked saccades

In several of the recording sessions the monkeys performed reliable, saccadic eye movements directed to the stimulation site within the retinotopic map in V1, namely the RFs of the stimulated neurons. This results is reproducing many previous studies of ICMS in V1 of alert and behaving monkeys (Bartlett et al. 2005; Bartlett and Doty 1980; Doty 1965; Ni and Maunsell 2010; Schiller et al. 2011; Tehovnik et al. 2003; Tehovnik and Slocum 2009). In contrast to these studies, where in most of them, the animals were trained to detect small visual targets and/or ICMS pulses, our animals were trained to maintain fixation (although in ICMS trials they were allowed to perform saccades within the fixation window). This fact can explain why the frequency of evoked saccades in our work was lower than previously reported, moreover, it is possible that the animals made saccades to the stimulated RFs despite their training paradigm (and not because they were trained to detect the appearance of target/phosphene/ICMS pulse). In addition, the stimulating electrode in our study was located mainly in the upper layers of V1, which were reported be less excitable for elicitation of saccades by ICMS (Bartlett et al., 2005; Tehovnik et al., 2003). There are few possible pathways through which ICMS can evoke saccadic eye movements, and we elaborate on two main possibilities. First, the neurons in the deep layers of V1 send direct projections to the superior colliculus (SC) (Fries 1984; Lund and Boothe 1975; Spatz et al. 1970; Swadlow 1983) which has a central role in saccade generation (Lee et al. 1988; Robinson 1972). The pyramidal cells in superficial V1 innervate those of deep V1 (Lund and Boothe 1975; Peters and Sethares 1991; Spatz et al. 1970) and it is also possible that current spread from upper layers activated axons that are directly projecting to the SC. Another option is that stimulation has caused the appearance of a phosphene in the monkey’s central visual field which attracted the animal attention, leading the animal to perform a saccade towards the stimulated location in the retnotopic map of V1. Finally, Schiller et al. (2011) claimed that the contrast of the evoked phosphenes varied from 2.6 to 10% implying that low contrast stimuli, especially at the detection threshold, will generate large variance in detection responses.

Assuming a V1 to SC origin for the ICMS evoked saccades, one would expect saccades with short latencies (80-120 ms; Peter H. Schiller, 1977; Stanford, Freedman, & Sparks, 1996; Tolias et al., 2005) and a small latency variance. The second explanation for ICMS evoked saccades suggests longer latencies and larger variability. The saccadic latencies we observed are in the range of 200 ms after ICMS onset and their large latency variance support the 2^nd^ explanation. However we cannot rule out the first possibility, because in our research the animals were trained to hold fixation and this may have interfered with saccades initiation, as previously reported (Tehovnik et al. 2004) (Tehovnik & Slocum, 2004).

Our results also show that trials with evoked saccades also show larger neuronal activity in response to ICMS. During normal visual processing, the stimulus intensity/saliency affects the response amplitude of the population response, for example stimuli of higher contrast will show larger VSD response (Meirovithz et al. 2010) (Meirovithz et al., 2010). It is possible to hypothesize that the opposite is also true – higher neuronal responses can reflect a more salient visual percept, even in cases where the perception is artificially created. This is in accordance with Schiller et al. (2011) and Winawer (2016) that reported that larger ICMS current or higher frequencies, can cause the appearance of phosphenes at higher contrasts. Thus it is possible that in trials where the monkey performed saccades to the RF of the stimulated area it was in response to the appearance of a more salient phosphene compared with trials that have shown a lower neuronal response.

